# Callose deposition is essential for the completion of cytokinesis in the unicellular alga, *Penium margaritaceum*

**DOI:** 10.1101/2020.05.28.122580

**Authors:** Destiny J. Davis, Minmin Wang, Iben Sørensen, Jocelyn K.C. Rose, David S. Domozych, Georgia Drakakaki

**Affiliations:** Department of Plant Sciences, University of California, Davis CA; Plant Biology Section, School of Integrative Plant Science, Cornell University, Ithaca, NY 14853; Department of Biology and Skidmore Microscopy Imaging Center, Skidmore College

**Keywords:** Callose, Cytokinesis, Penium margaritaceum, cell plate

## Abstract

Cytokinesis in land plants involves the formation of a cell plate that develops into the new cell wall. Callose is a β-1,3 glucan that transiently accumulates at later stages of cell plate development and is thought to stabilize the delicate membrane network of the cell plate as it expands. Cytokinetic callose deposition is currently considered specific to multicellular plant species as it has not been detected in unicellular algae. Here we present callose at the cytokinesis junction of the unicellular charophyte, *Penium margaritaceum*. Notably, callose deposition at the division plane of *P. margaritaceum* showed distinct, spatiotemporal patterns that could represent distinct roles of this polymer in cytokinesis and cell wall assembly. Pharmacological inhibition of cytokinetic callose deposition by Endosidin 7 treatment resulted in cytokinesis defects, consistent with the essential role for this polymer in *P. margaritaceum* cell division. Cell wall deposition and assembly at the isthmus zone was also affected by the absence of callose, demonstrating the dynamic nature of new wall assembly in *P. margaritaceum*. The identification of candidate callose synthase genes provides molecular evidence for callose biosynthesis in *P. margaritaceum*. The evolutionary implications of cytokinetic callose in this unicellular Zygnematopycean alga is discussed in the context of the conquest of land by plants.

**Summary Statement:** Evolutionarily conserved callose in *Penium margaritaceum* is essential for the completion of cytokinesis.

## Introduction

Cytokinesis features have long been used in comparative biology to infer evolutionary relationships (Balasubramanian et al., 2004; Miyagishima et al., 2008; Stewart et al., 1973). Even though it has been extensively studied in all major kingdoms of life, the precise mechanism(s) of cytokinesis and its change over evolutionary time remains one of cell biology’s biggest mysteries (Hall et al., 2008; Pollard, 2017). In the green plant lineage, cytokinetic characteristics have largely been studied in the context of understanding the migration of green plants from aquatic to terrestrial habitats and the cell wall modifications that occurred during this transition (Buschmann and Zachgo, 2016; McBride, 1967).

The land plant cell wall provides the plant with several key adaptations to terrestrial life including protection against abiotic stress and pathogen attack, structural support to the plant body, and the integration of environmental and developmental signals (Hall et al., 2008; Popper and Tuohy, 2010; Sørensen et al., 2010). *De novo* formation of the land plant cell wall occurs during cytokinesis as Golgi-derived vesicles carrying membrane and cell wall cargo coalesce at the plane of division to form a cell plate. The cell plate expands and matures centrifugally until it reaches the parental plasma membrane, thereby dividing the cytoplasm of the cell into two daughter cells (Drakakaki, 2015; Samuels et al., 1995; Seguí-Simarro et al., 2004; Smertenko et al., 2017). Callose, a β-1,3 glucan, is a vital luminal polysaccharide of the cell plate. This polymer is transiently incorporated into the cell plate where it is thought to act as a stabilizing matrix to the delicate membrane network while also aiding in the radial expansion of the cell plate as it matures into a cell wall (Jawaid et al., 2020; Samuels et al., 1995). Both genetic studies and pharmacological inhibition of callose deposition during cytokinesis have shown the essential role of callose in *Arabidopsis thaliana* cell plate maturation (Chen et al., 2009; Park et al., 2014; Thiele et al., 2009). Specifically, without callose, cell plate maturation stalls leading to fragmentation and failed cytokinesis, resulting in binucleate cells (Chen et al., 2009; Guseman et al., 2010; Park et al., 2014; Thiele et al., 2009).

The evolutionary significance of callose deposition at the cytokinetic junction has received little attention even though it is an essential component of cell division in land plants. Callose has been observed at the division plane of Streptophycean algae including *Chara* and *Coleochaete*, and filamentous/branched Chlorophycean algae (Scherp et al., 2001). Thus far, callose has only been reported at the cytokinesis junctions of *multicellular* plant species, leading to the hypothesis that callose is specifically associated with multicellularity in the green plant lineage (Scherp et al., 2001).

The lack of in-depth studies of cytokinetic callose in lineages closely related to land plants impedes our understanding of callose’s role in the evolution of land plant cytokinesis and cell wall formation. A valuable model to investigate this question is *Penium margaritaceum*, a unicellular freshwater desmid of the Zygnematophyceae; the group of streptophytes that are most closely related to land plants (Delwiche and Cooper, 2015). *P. margaritaceum* has a simple cylindrical shape and consists of two semi-cells, each containing a multi-lobed chloroplast that surrounds a nucleus at the cell center or the isthmus (Rydahl et al., 2014). *P. margaritaceum* has been valuable in cell biology studies as it is amenable to various live cell-based methodologies used in microscopy-based and experimental analyses (Rydahl et al., 2014). Immunological studies of *P. margaritaceum* have shown that its cell wall composition is notably similar to that of the primary walls of land plants (Domozych et al., 2014; Sørensen et al., 2011), making it an ideal system to study the evolution of land plant cell wall formation. *P. margaritaceum* contains a cellulosic inner layer and two outer pectic layers that include rhamnogalacturonan-I (RG-I) and a distinct homogalacturonan (HG) lattice that covers the semi-cells, with the notable exception of a narrow ring at the isthmus zone (Domozych et al., 2014), which is the site of wall polysaccharide deposition and assembly. The cell wall expands by the deposition of cellulose and pectin at the isthmus zone that displaces older cell wall material towards the poles of the cell. The isthmus is also the site of pectin demethylesterification with highly esterified HG being secreted first, followed by de-esterification and calcium binding to form the unique lattice (Domozych et al., 2014). The underlying cellular structure of the isthmus likely dictates the deposition and arrangement of new cell wall material. Identifying and characterizing events and structures at the isthmus zone is necessary to fully understand how *P. margaritaceum* grows and divides. By defining the events during *P. margaritaceum* cytokinesis we can begin to identify evolutionarily conserved cell wall features between later diverging land plants (i.e. embryophytes) and their aquatic ancestors.

In this study, we characterized cytokinesis in *P. margaritaceum* with a focus on callose deposition during cell wall formation. This provided novel insight into the evolution of the land plant cell wall and cytokinesis strategies of this alga and helped clarify our understanding of cell division mechanisms employed by unicellular and multicellular plants.

## Results

### ES7 treatment inhibits growth in P. margaritaceum cell cultures by disrupting cytokinesis

We took a pharmacological approach to dissect the role of callose in *P. margaritaceum* cytokinesis. In our earlier studies we identified and characterized Endosidin 7 (ES7), a small molecule that specifically inhibits callose deposition during cytokinesis in land plants (Drakakaki et al., 2011; Park et al., 2014). We tested the effects of ES7 on cytokinesis in *P. margaritaceum* and compared them with those previously observed in land plants. ES7 application inhibited *P. margaritaceum* growth in a concentration-dependent manner (Fig. 1A). Specifically, a reduction in cell culture optical density (a metric used to determine cell numbers) was observed upon treatment with ES7. The initial effect was first noticeable 4 days after ES7 application and was most apparent 7 days after treatment. In cultures supplemented with 5 µM ES7 for 7 days, cell numbers were reduced by ∼30% compared to the DMSO control, whereas cell numbers in cultures grown in 7.5 µM ES7 and 10 µM ES7 for 7 days decreased by ∼60% and ∼69%, respectively in comparison to the DMSO control (Fig. 1A). An initial delay in the inhibitory action of ES7 likely reflects an acclimation period required by cells to the new media containing ES7 prior to reaching the exponential growth phase. No significant differences were observed between *P. margaritaceum* cultures grown in growth media alone (NT) or in media containing the solvent (DMSO) employed to dissolve ES7 at any time point.

**Figure 1:**
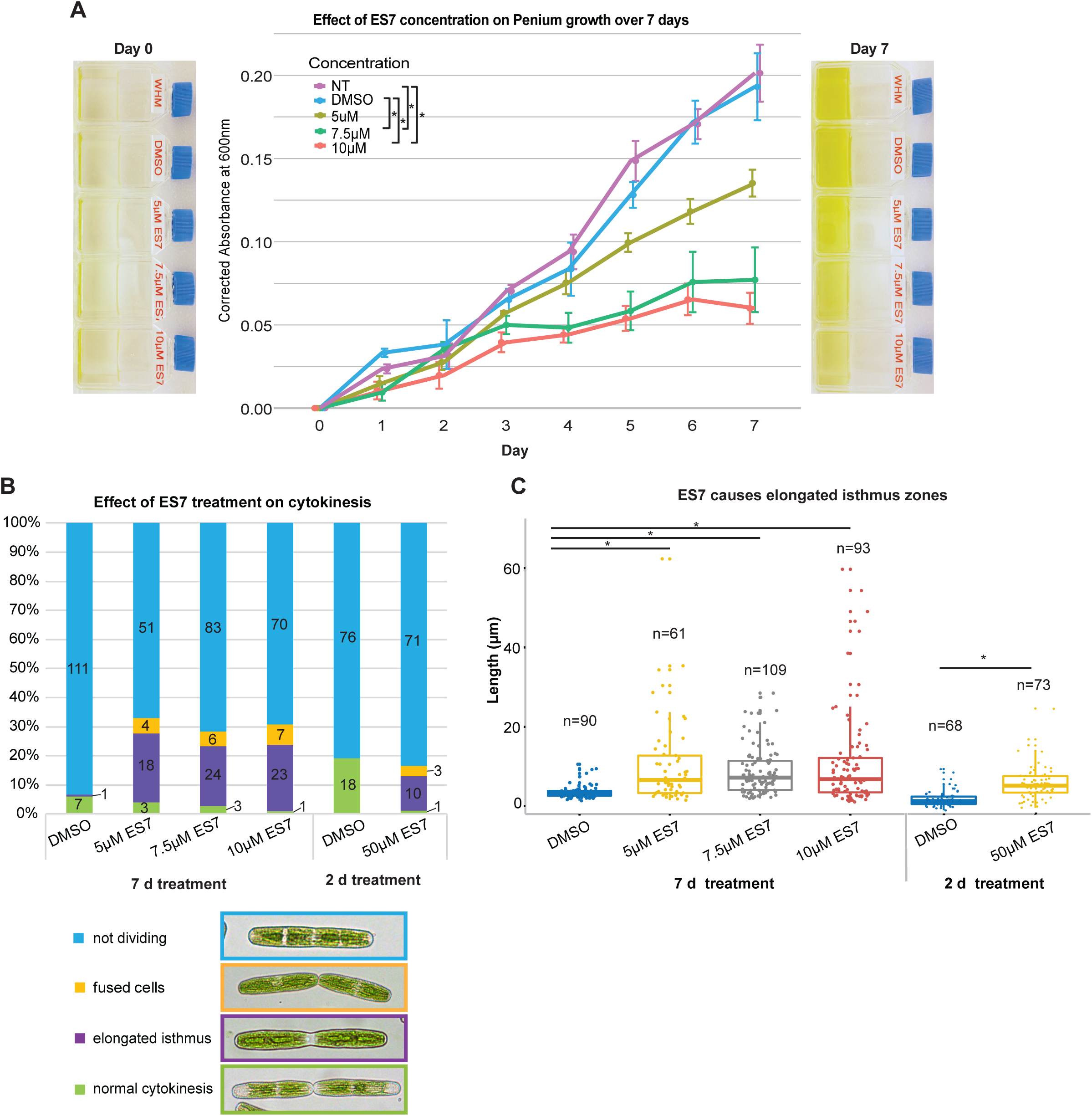
ES7 treatment inhibits cell proliferation by disrupting cytokinesis in a concentration dependent manner. (A) ES7 inhibition of *P. margaritaceum* liquid culture growth over a 7 day period measured by optical density (OD) at 600nm. Plot points represent the mean of 3 biological replicates, each measured 3 times per time point (technical replicates), with s.e.m. shown by the vertical lines. *P<0.05 measured by ANOVA with a Bonferroni correction (ns not shown for clarity). Representative images of day 0 and day 7 cultures are shown on each side of the graph. (B) Percentage of total cells exhibiting cytokinesis inhibition phenotypes in a series of low ES7 concentrations for 7 days and a 2 day high ES7 pulse treatment (cell count is centered in bars). Corresponding bright field images are shown below. (C) Box plot of isthmus zone measurements (µm) of cells in the same ES7 treatments as stated for (B). ***P<0.0005, 7 day treatment measured by Kruskal-Wallis rank sum test; 2 day treatment measured by Welch’s Two Samples t-test.

After noting the inhibitory effect of ES7 on cell culture growth, we then investigated whether this was due to inhibition of cytokinesis, as previously shown in ES7-treated *A. thaliana* seedlings (Park et al, 2014). We recorded the presence of cytokinesis phenotypes in the 7-day ES7-treated *P. margaritaceum* cultures and subsequently focused our attention on the isthmus zone (Fig. 1B). After 7 days of 5 µM, 7.5 µM, and 10 µM ES7 treatments we observed a higher proportion of fused cells (binucleate cells connected by a thin cytoplasmic strand, Fig. 1B yellow bars and yellow outlined brightfield image) and cells with atypically elongated isthmus zones when compared that of DMSO control cells (Fig. 1B purple bars and yellow outlined bright field image). We concluded that cells with elongated isthmus zones represented a recent inhibition by ES7 in early mitosis, whereas fused cells were those that had completed mitosis and had paused at cytokinesis due to the action of ES7. Fused cells always contained two daughter nuclei, indicating a defect in cytokinesis. In addition, fused cells were characterized by a thin cytoplasmic thread connecting two daughter cells, typically with a deep septum at the division plane (Fig. 1B: yellow outlined image). The fused cell phenotype was not observed in DMSO-treated cells but constituted 5.3% of the cells treated with 5 µM ES7, 5.2% of the cells in 7.5 µM ES7, and 6.9% of the cells in 10 µM ES7 (Fig. 1B). Non-dividing interphase (single nuclei) cells in control (DMSO) and ES7 treatments were indistinguishable. Control DMSO-treated cells exhibited elongated isthmus zones in only 0.8% of cells, versus 24% of cells grown in 5 µM ES7, 21% of cells grown in 7.5 µM ES7, and 23% of cells grown in 10 µM ES7 (Fig. 1B).

We then applied a short term (2 day), high concentration of ES7 (50 µM) to test the specificity of the inhibitor during cytokinesis in synchronized cultures and the reproducibility of the cytokinesis defects observed in 7 day, 10 µM ES7-treated cultures. Fused, binucleate cells that were unable to separate were observed in 12% of cells treated with 50 µM ES7 for 2 days compared to 0% of cells in DMSO (Fig. 1B). Cells with an elongated isthmus represented 3.5% of cells in ES7 compared to 0% of cells in DMSO (Fig 1B and Fig. 1C). The elongated isthmus of treated cells was approximately 3 times longer (averaged across all three ES7 concentrations) than the narrow isthmus found in control DMSO treated cells (Fig. 1C). We then carried out all subsequent experiments with cells treated with 50 µM ES7 for two days (∼48 hours) as this treatment allowed us to maintain a higher degree of cell cycle synchronization (Fig. 1B, green bars of 2 day DMSO compared to 7 day DMSO).

The cell phenotype and growth data showed that ES7 has a similar effect on dividing *P. margaritaceum* cells as on dividing *A. thaliana* root cells, i.e., daughter cells are unable to complete cytoplasmic separation following mitosis, resulting in binucleate cells (Park et al., 2014). Taken together, our results support an evolutionarily conserved ES7-inhibited pathway during cytokinesis between *P. margaritaceum* and land plants.

### Callose is present at the division plane of P. margaritaceum

Our previous studies have shown that ES7 causes cytokinesis defects in A thaliana by specifically inhibiting the deposition of callose at the cell plate (Park et al., 2014). Given the cytokinesis defect observed in ES7-treated *P. margaritaceum* cells, we hypothesized that callose plays a similar role in cytokinesis in this unicellular alga, challenging the view that callose is only present at the cytokinetic junctions of multicellular plant species (Scherp et al., 2001). Additionally, given the rapid progression of cytokinesis in *P. margaritaceum*, we posited that if callose is indeed present, it would be deposited during a very narrow time interval during late cell division, which in turn would make it difficult to observe. By closely examining the isthmus zone during cell division we were able to detect callose using two independent methods: a) aniline blue fluorochrome (Fig. 2, Evans et al., 1984); and b) with a monoclonal antibody with specificity for β-1,3 glucans (Fig. 3, Meikle et al., 1991). Callose was first apparent as a single ring at the isthmus zone of slightly elongated cells prior to mitosis (Fig. 2B), a likely indication of pending cell division (Ochs et al., 2014). We did not detect callose in interphase cells (Fig. 2A). Callose detection by aniline blue was more pronounced at the onset of cell division, with a bright aniline blue-labeled ring positioned at the isthmus zone during metaphase (Fig. 2C). The single ring was maintained as the daughter nuclei migrated to the isthmus zones of their respective daughter cells (Fig. 2D). As mitosis ended and cytokinesis began, the single callose ring transformed into two rings (Fig. 2E, 3D and surface rendering). As cell division progressed, the two callose rings (Fig. 2F arrowhead) surrounded a large aniline blue labeled punctum (Fig. 2F, arrow). Subsequently, the punctum expanded throughout the two adjoining ends of the daughter cells (Fig. 2G, arrow) while the two callose rings (Fig. 2G, arrowheads) began to disassemble. Finally, aniline blue labeling appeared as “caps” on the adjoining ends of the daughter cells (Fig. 2H, arrow) that remained for a period of time following the physical separation of the daughter cells (Fig. 2I). With these data we can summarize callose deposition into two general structures and associated stages: a) callose ring(s) characteristic of early cytokinesis events; and b) callose punctum/caps characteristic of later cytokinesis events (Fig. 2, arrowheads and arrows, respectively). We also observed these representative structures and stages in dividing cells labeled with the callose antibody (Fig. 3A and 3D). These results clearly demonstrate the presence of callose at the cytokinesis junction of a unicellular plant species, with a distinct deposition pattern in *P. margaritaceum*.

**Figure 2:**
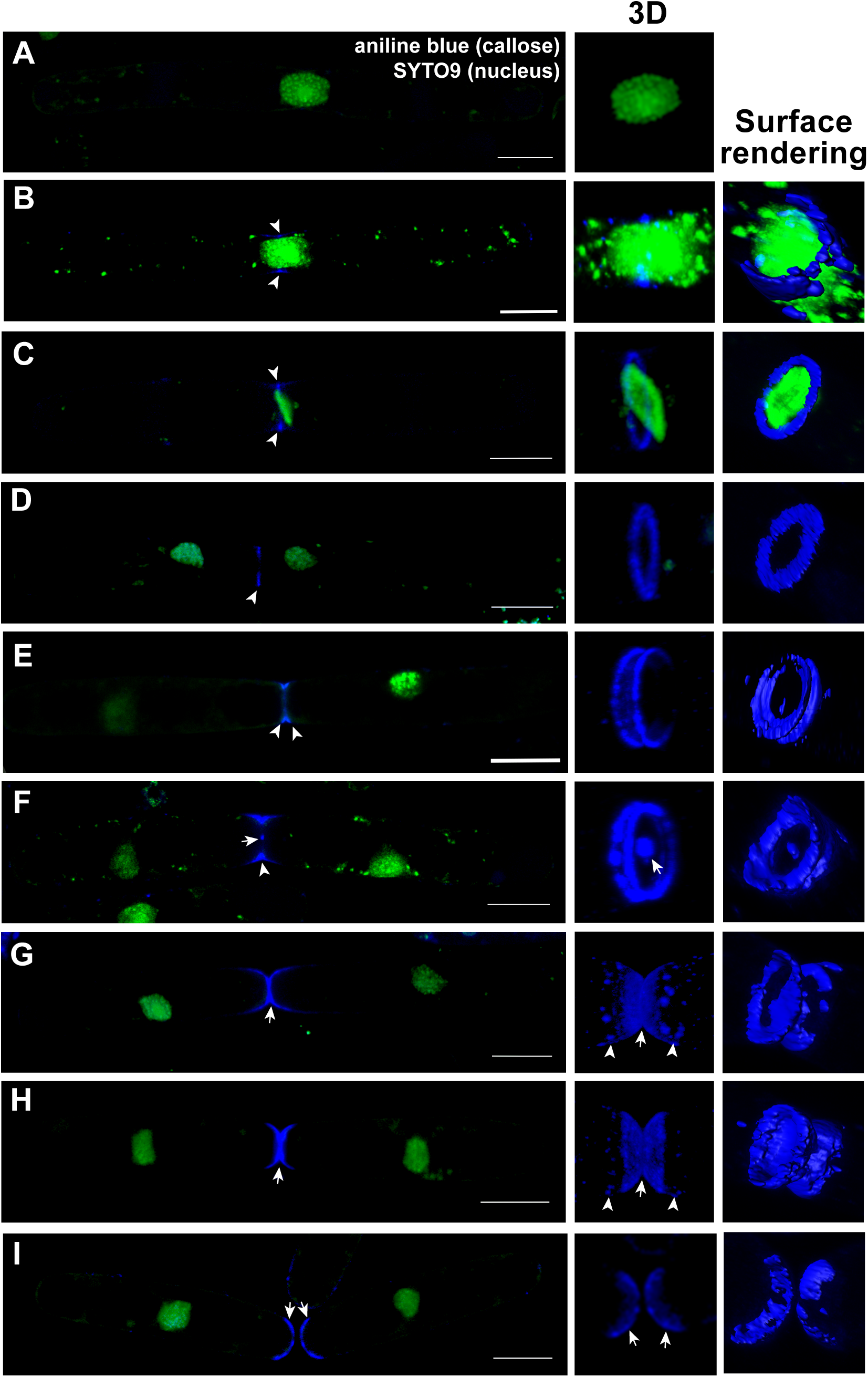
Callose is deposited in distinct patterns at the division plane of *P. margaritaceum*. Callose deposition patterns labelled by aniline blue fluorochrome during different stages of cytokinesis in *P. margaritaceum*. (A) Interphase cells did not show callose deposition at the isthmus zone (n = 26). (B) During the transition to mitosis a single aniline blue labeled ring was seen around the nucleus at the isthmus zone (n = 9) that remained during metaphase (C, arrowheads; n = 3) and persisted as daughter nuclei began migration to their respective isthmus zones (D, arrowheads; n = 15). When daughter nuclei reached their isthmus zones, callose was observed as two rings (E, arrowheads; n = 12). At the next stage, a bright aniline blue labeled punctum (F, arrow) was present in the center of the two rings (F, arrowheads; n = 5). During the ring disassembly stage (G, arrowheads), the punctum expanded and the isthmus zone constricted to form the daughter cell tips (G, arrow; n = 9). The next stage is characterized by opposing callose caps at the poles of the dividing cells and the disappearance of the callose rings (H, arrows; n =18). Callose caps remained until the daughter cells physically separated (I arrows; n = 26). 3D images (center column) produced by Zeiss Zen Black imaging software. Surface renderings shown to the right produced with Imaris (Bitplane). Nucleus labeled by SYTO9 are shown in green; callose labelled by aniline blue fluorochrome is shown in blue. Scale bars = 20 µm.

**Figure 3:**
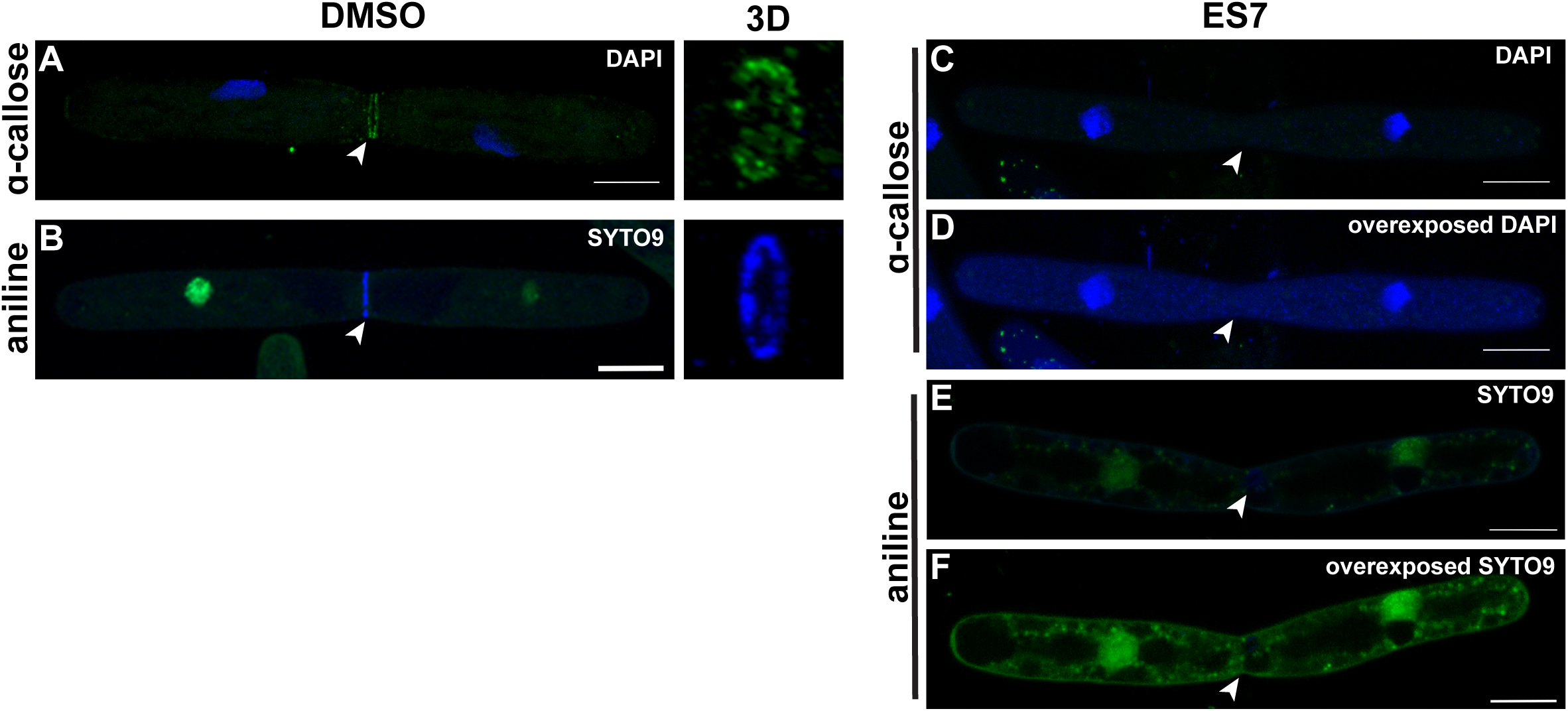
Inhibition of callose at the isthmus zone leads to cytokinetic defects in *P. margaritaceum*. Immunostaining of callose at the isthmus zone of dividing cells verify the presence of the polymer (A; n = 15). Immunolabeling of callose mimicked the pattern observed with aniline blue fluorochrome (A and B). ES7 treatment inhibits callose deposition (C-F). The polymer is no longer detectable by callose antibodies (C and D; n = 9) or aniline blue (E and F; n = 20) at the isthmus zone of fused daughter cells following ES7 treatment, visualized by overexposed fluorescent nuclear labeling (D and F). Scale bars = 20 µm.

### Callose deposition is essential for cytokinesis in P. margaritaceum

Given that ES7 arrested cell division in *P. margaritaceum* we investigated whether, like in A. thaliana, it inhibited callose deposition at the division plane. In contrast to the callose rings observed in DMSO-treated control cells during early cytokinesis (Fig. 3A and B, arrowheads and 3D insets), no callose was detected either immunologically or by aniline blue staining in dividing cells treated with ES7 (Fig. 3C-F), mimicking the effect in dividing *A. thaliana* root tip cells (Park et al., 2014). This indicates that callose is required for the progression of cytokinesis and suggests an evolutionarily conserved role for β-1,3 glucans during cell division among unicellular streptophyte algae and land plants.

### Callose deposition is not essential for septum formation

A key difference between *P. margaritaceum* and land plants is the presence of a septum during cytokinesis (Pickett-Heaps et al., 1999). The *P. margaritaceum* septum manifests itself primarily as a curvature of the cell wall and constriction of the plasma membrane during the physical separation of daughter cells (Fig. 4A, arrowheads). Septum formation occurs at the isthmus and coincides with callose deposition (Fig. 4A and B). We sought to determine if ES7 inhibits cell separation by disrupting septation. ES7 treatment did not alter the formation of the cell wall curvature and/or plasma membrane constriction that constitute the septum (Fig. 4E arrowheads), even as callose was no longer detected (Fig. 4F), indicating that callose does not play a major role in septation.

**Figure 4:**
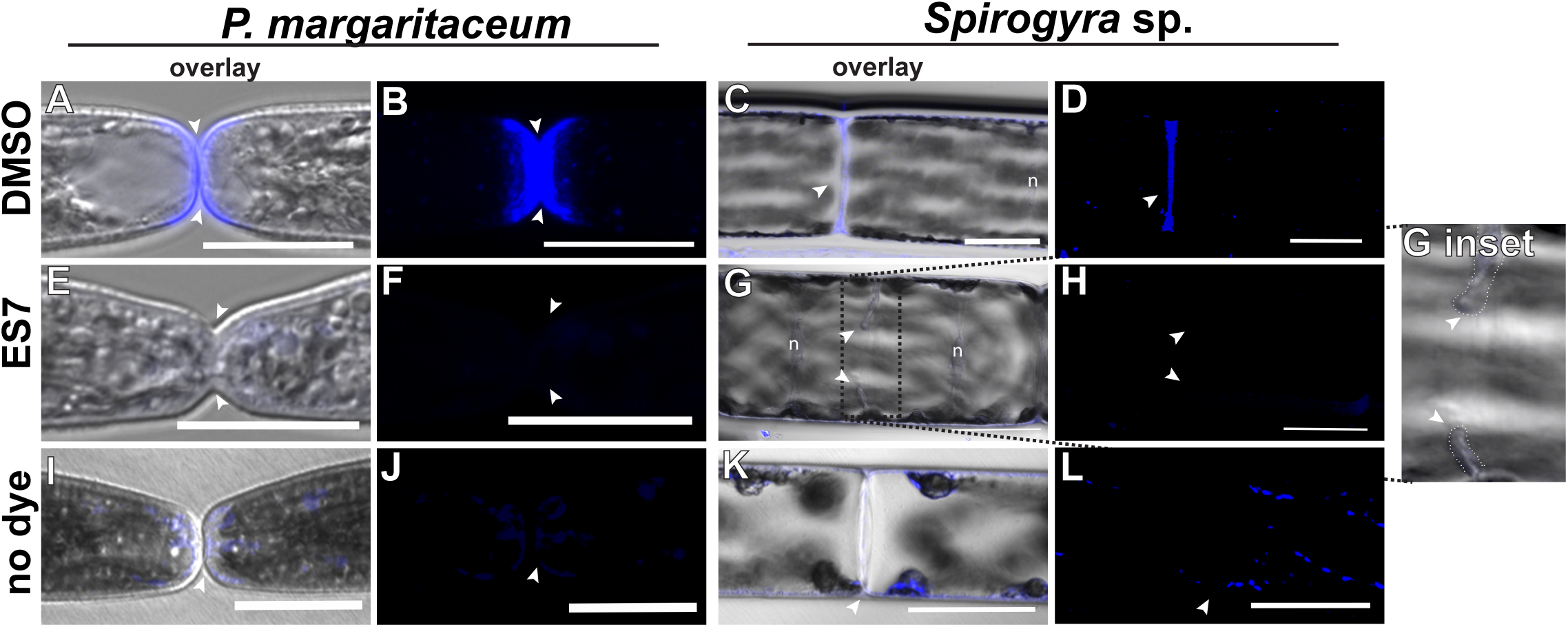
ES7 treatment does not inhibit septum formation. Callose was detected by aniline blue fluorochrome (blue) at the division plane of *P. margaritaceum* (A, B) and *Spirogyra* sp. (C, D; n = 39), coinciding with septum formation (arrowheads). Following ES7 treatment, callose was no longer detectable at the division plane of either species (F and H); however, septa characterized by isthmus zone constriction in *P. margaritaceum* (E, arrowheads) and membrane ingrowths in *Spirogyra* sp. (G, arrowheads; n = 5), were still visible in both species. Enlarged inset of (G) showing septum stubs (dashed outline) despite the absence of callose shown to the far right. Dark field images are max intensity projections. *P. margaritaceum* scale bars = 20 µm; *Spirogyra* sp. scale bars = 50 µm. n = nucleus.

The rapid series of events that occur at the division plane of *P. margaritaceum* makes distinguishing septation from other cytokinesis components, including a potential cell plate, very difficult. Cytokinesis in the closely related multicellular zygnematophyte, *Spirogyra* sp., however, has been well characterized, including the processes of septation and cell plate formation (Fowke and Pickett-Heaps, 1969; Sawitzky and Grolig, 1995). Taking advantage of this, we analyzed dividing *Spirogyra* sp. cells for the presence of callose and for an ES7 effect on cell division. *Spirogyra* sp. also deposited callose at the plane of division (Fig. 4C and D). Treatment of *Spirogyra* sp. cells with ES7 resulted in loss of callose, determined by the absence of aniline blue labeling and disruption of cytokinesis observed by the occurrence of binucleate cells and septum ingrowths (Fig. 4G and inset). Assuming that the curvature of the adjoining *P. margaritaceum* daughter cells is analogous to the septum in *Spirogyra* sp., our data suggest that ES7 does not significantly affect septum formation, but rather perturbs a callose-dependent process during cytokinesis, and in the case of *Spirogyra* sp, cell plate formation specifically.

### Lack of callose alters the deposition pattern of polysaccharides during cell wall formation

The major polysaccharides in the mature cell wall of *P. margaritaceum* are cellulose and pectins, which are deposited at the isthmus and displace older cell wall components towards the two poles (Domozych, 2014; Domozych et al., 2007). As the isthmus is the focal point of both callose deposition and the deposition of wall polymers, we assessed the impact of the lack of callose, due to ES7 treatment, on subsequent cell wall development. During interphase in both control and ES7 treated cells, low methyl esterified homogalacturonan (HG) labeled by the monoclonal antibody (mAb), JIM5, showed a typical lattice pattern across the entire cell wall surface, with the exception of a narrow unlabeled band at the isthmus zone (Fig. 5A and B; E and F). As the cells elongated prior to division, the narrow unlabeled isthmus zone expanded to a wider band (Fig. 5C and D). However, in cells arrested in cytokinesis by ES7, JIM5 labeling was seen covering the isthmus zone between two fused daughter cells (Fig. 5G and H, inset). While the low methyl esterified HG in the older areas of the cell wall (i.e. away from the isthmus zone) maintained a normal lattice-like appearance (Fig. 5H inset, arrow), JIM5 labeling of the elongated isthmus produced a weakly fluorescent signal with no lattice, but instead a rather irregular pattern (Fig. 5H and inset, arrowheads).

**Figure 5:**
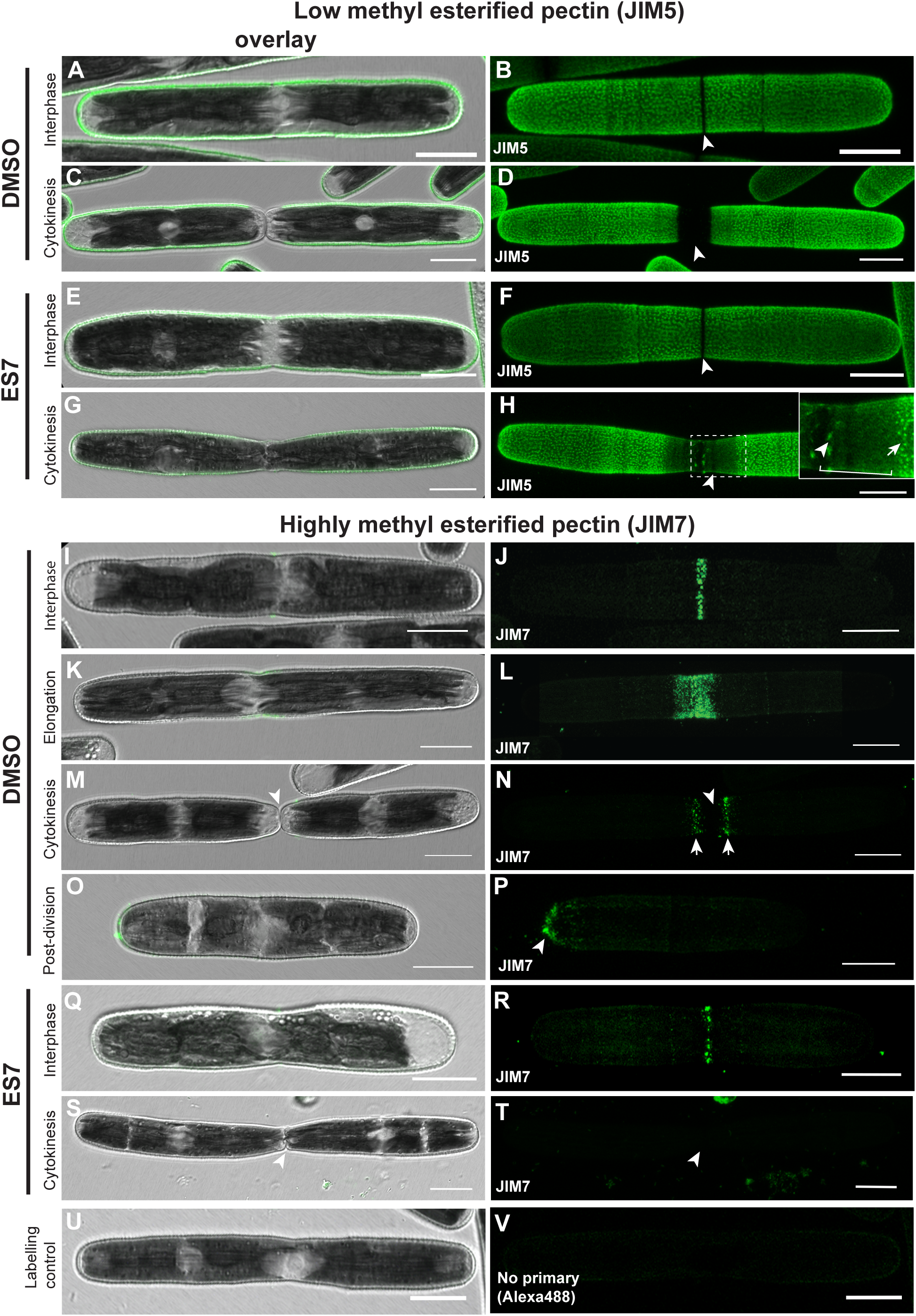
Pectin patterning changes in response to callose inhibition by ES7 treatment. (A and B) In DMSO control cells during interphase, low methyl esterified homogalacturonan (HG) labeled by JIM5 covered the entire cell surface in a regular lattice pattern, except for a narrow unlabeled ring at the isthmus zone (arrowhead; n = 76). (C and D) During cytokinesis, the unlabeled isthmus ring from (B) expanded to a wide, unlabeled band (arrowhead; n =21). (E and F) ES7 treated interphase cells were indistinguishable from DMSO control (n = 54). Dividing cells treated with ES7, however, exhibited JIM5-labeled HG punctum in the isthmus zone (G, H arrowheads; n = 23). Notably, the normal JIM5 lattice pattern (H inset, arrow) was not maintained in the wall at the ES7 elongated isthmus (H inset, arrowheads). In contrast to JIM5 labeled HG, highly methyl-esterified HG labeled by JIM7 antibodies was typically observed as a single narrow band at the isthmus zone (I and J) during interphase (n = 13). During cell elongation prior to division, the JIM7 labeled HG band expanded (K and L; n = 12) and eventually formed two distinct bands (arrows) on each side of the division plane (M and N, division plane marked by arrowhead; n = 9). (O and P) Upon immediate physical separation of the daughter cells, JIM7 labeling remained at the new cell poles (arrowhead; n = 4). (Q and R) JIM7 labeled HG in ES7 treated interphase cells was indistinguishable from DMSO control (n = 6). (S and T) During cytokinesis, however, JIM7 labeling was completely absent (isthmus of fused cell indicated by arrowhead; n = 21). Staining control (no primary antibodies, Alexa488 only) shown in U and V. Fluorescent images are max intensity projections. Scale bars = 20 µm.

High methyl esterified HG labeled by the mAb, JIM7, was typically found exclusively at the isthmus zone (Fig. 5I and J), consistent with previous observations (Domozych et al., 2011). Upon elongation, control treated cells exhibited a wide JIM7-labeled HG band at the isthmus zone (Fig. 5K and L), which later transformed into two distinct JIM7-labeled bands on either side of the division plane (Fig. 5M and N, arrows). Upon physical separation, the daughter cells retained JIM7 labeling at the once-connected cell poles, indicating continued cell wall maturation at the new cell poles (Fig. 5O and P). ES7 treated interphase cells showed no difference in JIM7 labeling compared to DMSO treated control cells (Fig. 5Q and R). However, JIM7 labeling was absent from the isthmus zones of ES7 treated cells that exhibited a fused cell phenotype (Fig. 5S and T).

To gain a clearer view of cell wall dynamics and cytokinesis events during and following ES7 treatment, we combined cell wall labeling (JIM5) with ES7 washout, thereby removing ES7 by washing cells with DMSO supplemented Woods Hole Medium (WHM), and observed cell wall growth and cytokinesis recovery. Following 48 hours of recovery in fresh growth media, cells exhibited normal JIM5-labeled HG deposition patterns: lattice over the semi cells with a narrow unlabeled isthmus zone that expanded to a large band during elongation (Fig. S1A-D). Comparing initial cell wall labeling (green) to new cell wall growth in recovery conditions (magenta) revealed that the majority of cells (90.1%) elongated and/or divided normally (Fig. S1A-D). Some cells, however, retained ectopic JIM5-labeled HG in the isthmus zone and were devoid of new pectin deposition at the daughter cell isthmus zones where elongation would occur (Fig. S1E), indicating no growth in recovery conditions. One cell, however, deposited new cell wall both at the fused parental isthmus zone and the daughter cell isthmus zones, demonstrating cell wall growth despite the cytokinesis arrest (Fig. S1F). The presence of fused cells following recovery suggests that the deposition of callose during cytokinesis is a significant checkpoint in *P. margaritaceum* cell division, from which it never recovers once interrupted.

Cellulose labelling with a carbohydrate binding module specific to crystalline cellulose fused to green fluorescent protein (GFP-CBM3a) was observed as either one or two bright, narrow rings at the isthmus zone, consistent with previous observations (Fig. S2A-C, Domozych, 2014). The difference in signal intensities between the mature cell wall and the isthmus is likely due to easier access of the probe to the isthmus zone, since this region of the cell wall is devoid of mature pectin (Domozych, 2014). While the typical GFP-CBM3a labeled single ring was seen in ES7-treated fused cells (Fig. S2D and E), we also observed fused cells with multiple rings off-center from the isthmus zone (Fig. S2F), within the isthmus zone (Fig. S2G), or multiple rings near the isthmus zone (Fig. S2H).

In summary, cell wall polymer labelling showed that a lack of callose in the isthmus zone of dividing cells and the subsequent arrest of cytokinesis affects the deposition pattern of both cellulose and pectin specifically at the isthmus zone during cytokinesis. Additionally, these effects do not induce changes in the older cell wall. Our data further show that inhibition of cytokinetic callose does not affect the biosynthesis of structural polysaccharides, but likely influences the architecture of the isthmus zone, which may indirectly affect the polysaccharide deposition pattern.

### Glucan synthases-like (GSL) genes of P. margaritaceum are more closely related to those of land plants than to homologs in the Chlorophyta

Glucan synthases-like (GSL) proteins are enzymes that incorporate UDP-glucose into β-1,3 glucan and are responsible for callose deposition at the division plate in higher plants (Chen et al., 2009; Verma and Hong, 2001). Analysis of representative plant GSL proteins ranging from chlorophyte algae to *A. thaliana* by NCBI conserved domain search (Lu et al., 2019) revealed them to be large proteins of ∼2,000 amino acids, consisting of the glucan synthase domain and the glucan synthase subunit FKS1 domain. In addition, the regulatory domain Vta1 is prevalent in land plant GSL proteins near the N-terminus (Fig. S3).

The similar ES7-based inhibition of cell division and callose deposition between *P. margaritaceum* and *A. thaliana* suggests that their respective GSL proteins have similar molecular properties. To identify candidate *P. margaritaceum* proteins we searched the newly released *P. margaritaceum* genome assembly and transcriptome (Jiao et al., 2020) using BLAST analysis (Altschul et al., 1990) with the 12 *A. thaliana* GSL isoform sequences as the query. This revealed 15 putative full-length *P. margaritaceum* GSL proteins, four of which showed > 50% amino acid similarity compared with AtGSL2, an *A. thaliana* GSL with the glucan synthase domain, FKS1 domain and the regulatory Vta1 domain (Fig. S3). The other eleven *P. margaritaceum* sequences showed < 30% amino acid similarity against AtGSL2 (Fig. S3). The four *P. margaritaceum* GSLs that showed >50% amino acid similarity against AtGSL2 were used in subsequent phylogenetic analyses.

A phylogenetic analysis of full-length GSL proteins from algae and land plants revealed that chlorophyte algae form an outgroup in an unrooted tree (Fig. 6, blue arrow), while *P. margaritaceum* and *Klebsormidium flaccidum*, a multicellular charophyte, are more closely related to the GSL proteins of land plants (Fig. 6, black arrow). In the clade containing pm0123640g0030, pm000009g0010, and pm010253g0010, all the land plants have at least one copy of a GSL homolog, suggesting that all the land plants have homologs of ancestral charophyte GSL genes (Fig. 6, black arrow). Phylogenetic analysis of plant GSL proteins from the Chlorophyta, Charophyta and Embryophyta suggested that *P. margaritaceum* GSL proteins are more closely related to those from land plants than to those from chlorophyte algae.

**Figure 6.**
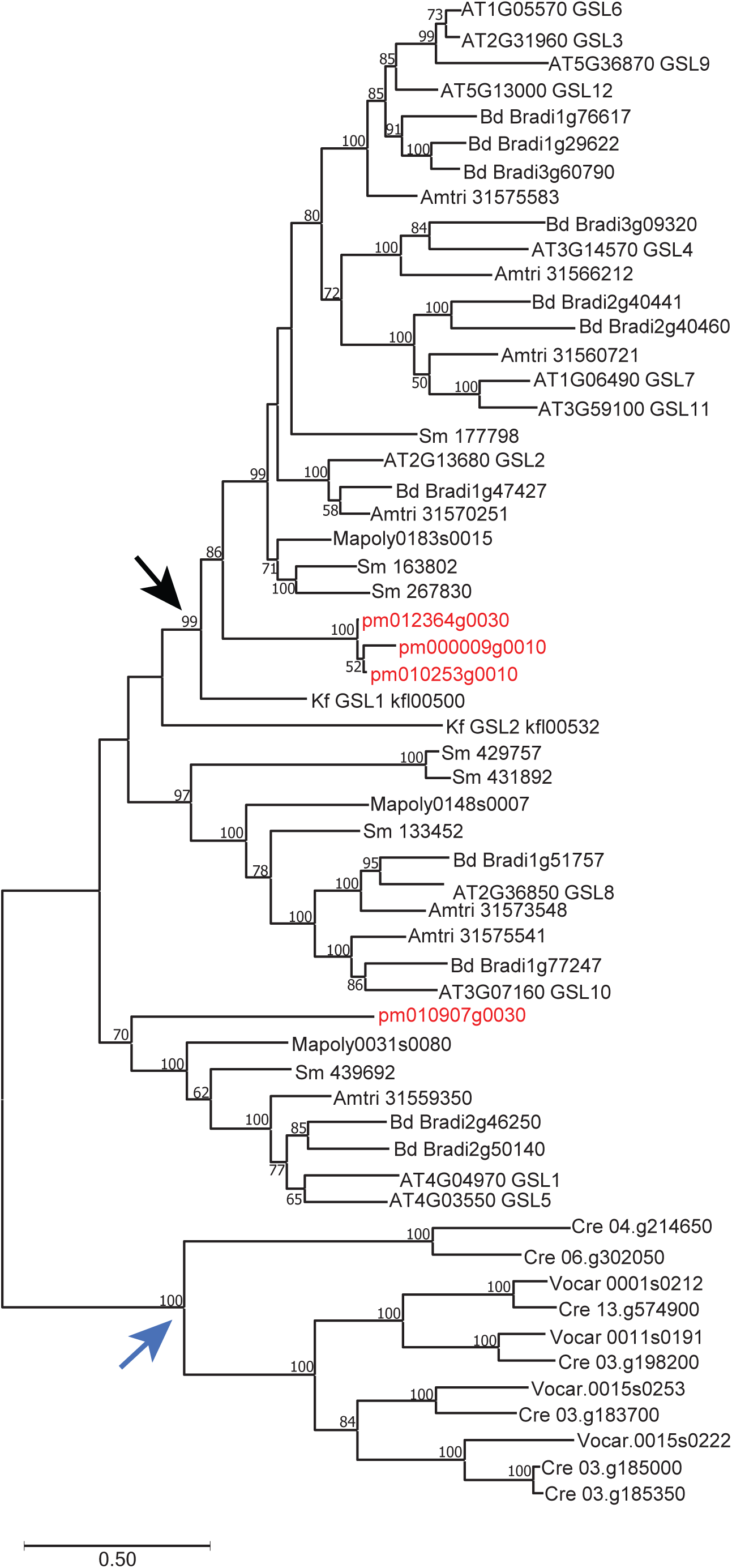
Phylogenetic analysis of glucan synthase-like enzymes. GSLs of representative species of Embryophyta, Charophyta, and Chlorophyta are aligned by T-COFFEE and analyzed in MEGA to construct an unrooted maximum-likelihood tree of *P. margaritaceum* and other plants. Bootstrap values are labeled at each branch (branches with a bootstrap value below 50 are left as unresolved) and evolutionary distance is proportionally drawn by branch length. Phytozome accession numbers were used to name the enzymes, except for *P. margaritaceum* sequences which were obtained from Jiao et al. 2020 and *Klebsormidium flaccidum* sequences that were obtained from the *Klebsormidium* genome project website. Amtri, *Amborella trichopoda*; At, *Arabidopsis thaliana;* Bd, *Brachypodium distachyon;* Cre, *Chlamydomonas reinhardtii;* Kf, *Klebsormidium* flaccidum; Mapoly, *Marchantia polymorpha;* Pm, *Penium margaritaceum*; Sm, *Selaginella moellendorffii*; Vocar, *Volvox carteri*.

## Discussion

### Callose is deposited at the division plane of P. margaritaceum and is necessary for the completion of cytokinesis

Callose deposition in land plants is an essential component of cell wall formation during cytokinesis (Samuels et al., 1995). Without the tight temporal and spatial deposition of callose at the expanding cell plate, the transition of the cell plate to a nascent cell wall is arrested and daughter cells fail to separate, leading to binucleate cells (Chen et al., 2009; Guseman et al., 2010; Park et al., 2014; Thiele et al., 2009). It has been proposed that only multicellular species of plants take advantage of the unique elastic properties of callose during cytokinesis (Scherp et al., 2001). Given the wide variety of plant body phenotypes and a diverse array of transitional cytokinetic characteristics in the algal lineages most closely related to land plants, a key question remains as to whether callose deposition during cytokinesis is strictly a multicellular characteristic (Hall et al., 2008).

Our earlier studies established ES7 as a highly specific tool in dissecting the role of callose during cytokinesis in *A. thaliana* (Drakakaki et al., 2011; Park et al., 2014). Treatment of *P. margaritaceum* with ES7 has a similar cytokinesis effect as in A. thaliana, leading to the formation of binucleate cells (Park et al., 2014). In land plants, it is hypothesized that callose plays an essential role in providing mechanical support of the cell plate and aids in its expansion (Jawaid et al., 2020; Samuels et al., 1995). While a cell plate was not visible in our light microscopy experiments with *P. margaritaceum*, the clear inhibition of cytokinesis following ES7 treatment suggests that callose also has an essential function in its cell division. Many details of the role(s) that callose plays in the cell division of *P. margaritaceum* remain to be determined; however, the distinct temporal and spatial callose patterning and ES7 effects provide valuable insights. For example, the highly elongated isthmus zone of ES7-treated pre-mitotic cells may reflect a role for callose in demarking the narrow area of the isthmus zone by maintaining the underlying isthmus zone structure. The occasional presence of a callose ring at the isthmus zone of pre-mitotic cells (Fig. 2B) could support this hypothesis. It is unclear whether a callose ring is a constant fixture of the isthmus zone or if it appears only in preparation for cytokinesis, given that it is not seen in every pre-mitotic cell.

### Distinct callose patterns characterize cytokinesis stages in P. margaritaceum

It is not until the onset of cytokinesis that callose is seen as temporally distinct, brightly labeled patterns by aniline blue staining: first, aniline blue-labeled rings and later aniline blue-labeled caps. Ringed callose always preceded the presence of callose caps and is the only callose pattern seen prior to daughter cell nuclei migration. Because of this temporal separation, we can define early cytokinetic events as those with ringed callose and late cytokinetic events as having callose caps. The transition between these two patterns and stages of cytokinesis is also clearly distinguished with the appearance of the conspicuous central callose punctum between the two callose rings (Fig. 2E). In subsequent stages the callose foci appear to expand to form the caps while the two callose rings disassemble, visualized as aniline blue labeled fragments (Fig. 2F). We propose that the distinct ring and cap callose deposition patterns have distinct functions but are both required for proper cell separation. Callose hydrogels, with their ability to associate with cellulose to reduce its rigidity (Abou-Saleh et al., 2018), are likely advantageous in covering the dome-shaped apex of dividing cells. Specifically, the elastic properties of callose hydrogels could provide a flexible barrier during separation while maintaining a curved shape at the new daughter cell poles. In addition, the callose dome itself may be the initial loadbearing support for the cell wall and provide a mechanism to overcome tension stress, as has been previously proposed for pollen tube growth (Parre and Geitmann, 2005). Evidence from aeroterrestrial algae such as *Klebsormidium* point towards a mechanism of callose deposition in shoring up areas of the cell wall prone to experiencing biomechanical stress due to loss of turgor pressure (Herburger and Holzinger, 2015). In this model, the elastic properties of callose enhance cell wall flexibility to a degree that is necessary for life in habitats with fluctuating water availability; a feature thought to be key for dispersal and led to the widespread success of these algae (Holzinger et al., 2015). The increased deposition of callose during water scarcity paired with the lack of typical load-bearing components of the cell wall and glycoproteins in *Klebsormidium* suggest that callose deposition may be an ancient mechanism for providing cell wall support in a variety of habitats and developmental stages.

Cytokinesis in *Spirogyra* sp. also occurs in distinct phases, as shown by chemical inhibition and physical disruption of cytoskeletal structures at specific time points (McIntosh et al., 1995). Specifically, depolymerization of phragmoplast microtubules was reported to inhibit complete cross-wall closure, while the septum was not affected, indicating that septation alone is not sufficient for the completion of cytokinesis (McIntosh et al., 1995; Sawitzky and Grolig, 1995). Our experiments here with ES7 treatments of *Spirogyra* sp. point to the same conclusion, in that inhibition of cell plate formation resulted in cytokinesis defects but had no effect on septum initiation (Fig. 4G and inset). Collectively our data, consistent with findings for other Zygnematophycean species (Ochs et al., 2014; Sawitzky and Grolig, 1995), suggest that cytokinesis in *P. margaritaceum* represents a transitional form of cell division that emerged prior to the first land plants. This hypothesis is based on several key pieces of evidence: 1) callose is essential for the completion of cytokinesis in land plants and both algae species observed in this study, and specifically for cell plate formation in *Spirogyra* sp. and A. thaliana; 2) In *P. margaritaceum*, the central callose punctum appears at the last physical location of daughter cell connection, where a cell plate is expected; and 3) the central callose punctum appears to expand centrifugally, as in *Spirogyra* sp. and land plants, in contrast to the centripetal growth of the septum.

The presence of the callose rings is intriguing. In fission yeast, enriched β-1,3 glucan deposition characterizes septum formation and is regulated by the activity of specific glucan synthases (Cortés et al., 2016). In dividing yeast cells, callose is deposited in the secondary septum once the chitin-rich primary septum (PS) has developed (Fraschini, 2020). Our study showed that septum formation is still achieved in the absence of callose, as evidenced by the clear furrows of ES7-treated *Spirogyra* sp. and *P. margaritaceum* (Fig. 4). Therefore, we hypothesize that callose aids in supporting the membrane as it grows inward during new daughter cell pole formation in cytokinesis but is not required for septum initiation. The highly elongated and fused isthmus zones of ES7-treated cells is consistent with this idea. A role of callose in supporting septum formation between yeast and *P. margaritaceum* could indicate evolutionary conservation across eukaryotes.

### Callose inhibition during cytokinesis may affect cell wall polysaccharide assembly by perturbing isthmus zone architecture

The isthmus zone of *P. margaritaceum* is a highly dynamic area of the cell wall. Not only is it the location of division, but it also marks the area of new cell wall deposition and assembly at all stages of the cell cycle (Domozych et al., 2011). Cell wall growth regarding pectin is follows a consistent sequential pattern beginning with the deposition of highly esterified HG, followed by cell wall displacement, de-esterification and calcium crosslinking. We hypothesize that highly esterified HG is deposited at the isthmus zone *and*, only in the final stages of cytokinesis, at a location interior to the isthmus zone growing outward centrifugally, as a cell plate would grow, to eventually cover the daughter cell poles once separated. As stated previously, we hypothesis the central callose punctum marks the presumed cell plate in dividing *P. margaritaceum* cells. However, further experimentation involving high resolution microscopy is necessary to identify the callose punctum as a cell plate and elucidate polysaccharide deposition dynamics interior to the isthmus zone. Whether and in what form pectin deposition occurs at the callose punctum is of particular interest given the role pectin play in cell adhesion of plant cells (Daher and Braybrook, 2015).

It was apparent from our studies that while the older cell wall was not affected by ES7-mediated cytokinesis inhibition, the wall at the isthmus zone was more susceptible to changes. Dividing cells treated with ES7 could compensate for the lack of callose by either synthesizing an excess, or directing the deposition of, polysaccharides such as cellulose and pectin to the isthmus zone (Fig. 5H, inset and Fig. S2F-H). The loss of the normal JIM5 lattice pattern together with the presence of weak and or aberrant JIM5 labeling in the isthmus may represent an effect on the underlying structure of the isthmus zone during cytokinesis, as a consequence of the loss of callose following chemical treatment. It is plausible that the lack of callose changes some conformational component at the isthmus so that the deposition pattern of HG is disrupted (Domozych et al., 2014).

Additionally, increases in HG deposition with a low level of esterification paired with increases in calcium crosslinking to compensate for the loss of another cell wall polysaccharide, such as cellulose, is a common feature of cell wall plasticity (Díaz-Cacho et al., 1999; Manfield et al., 2004; Sabba et al., 1999; Shedletzky et al., 1990). The absence of callose during cytokinesis may lead to the deposition of the predominant wall polymer in *P. margaritaceum*, namely partially esterified HG (Domozych et al., 2007), strengthening linkages with calcium ions and stabilizing the isthmus zone. This is probably achieved by an increase in pectin methylesterase activity at the isthmus zone in response to the inability to separate. The lack of highly methylesterified HG labeled by JIM7 in ES7-treated cells points to a similar conclusion (Fig. 5T). Our recovery experiments also revealed cells that were unable to divide even after ES7 washout (Fig. S1E and F), indicating the presence of a checkpoint that, when disrupted, prevents the cell from progressing through cytokinesis.

Notably, we saw no obvious effect on any of the tested cell wall polysaccharides other than at or near the isthmus zone of dividing cells following ES7 treatment, highlighting the specific action of ES7 and potentially the role of callose in *P. margaritaceum*. In addition, polysaccharide deposition is always maintained in a ring-shaped pattern, suggesting that targeted secretion mechanisms of polysaccharides are likely not altered by ES7 treatment, consistent with our observations in *A. thaliana* (Drakakaki et al., 2011; Park et al., 2014).

### Evolutionary insights of cytokinetic callose and putative callose synthase proteins in P. margaritaceum

Callose is involved in many physiological processes, including response to wounding, regulation of intercellular transport through plasmodesmata, pollen tube growth, cell wall reformation in protoplasts, and cell plate formation (Piršelová and Matušíková, 2013). Interestingly, wound-induced callose deposition have been observed across evolutionarily distinct taxa including unicellular chlorophyte algae, filamentous streptophyte algae and land plants (Jacobs et al., 2003; Scherp et al., 2001; Stone and Clarke, 1992). This is consistent with the presence of a number of callose synthases in the genomes of a wide range of plant species from chlorophytes to embryophytes (Fig. 6, Drábková and Honys, 2017; Ulvskov et al., 2013; Verma and Hong, 2001). The presence of callose at division planes appears to be a more recent evolutionary feature, having only been seen in multicellular species. The presence of callose in *P. margaritaceum* likely reflects an evolutionary remnant from an ancestor that utilized callose in cytokinesis. The unicellular and filamentous Zygnematophyceae are thought to be derived from a morphologically complex ancestor (de Vries and Archibald, 2018; Delwiche and Cooper, 2015). The remarkable morphological reduction paired with the wide diversity of cell division patterns in the Zygnematophyceae probably aided in the evolution of terrestrial plant life in that simpler body plans are better able to survive rapidly changing habitats and variable water availability (Delwiche and Cooper, 2015; Hall et al., 2008).

Analysis of the *P. margaritaceum* genome (Jiao et al., 2020) revealed the presence of putative callose synthases. Phylogenetic analysis indicated that they have a closer relationship to homologs from land plants than to those from chlorophytes. The identification of putative callose synthases in *P. margaritaceum* supports the presence of callose as demonstrated by the immunodetection and pharmacological inhibition of this study. However, the *P. margaritaceum* isoform(s) responsible for callose deposition at the isthmus zone have yet to be determined by complementation using *A. thaliana* mutants—an important next step in our future work with *P. margaritaceum*.

Based on our results, we present a model of cytokinesis in *P. margaritaceum* (Fig. 7) that includes an essential role for callose in the final stages of cell division. This represents an important advance in our understanding of the evolutionary significance of cytokinetic callose—one that includes a unicellular plant species. Whether *P. margaritaceum* is an isolated example of a unicellular algae using callose during cytokinesis, or if other examples among the unicellular streptophyte algae exist remains to be determined. Detailed studies identifying the role(s) of cytokinetic callose in early diverging plant lineages are crucial to fully grasp the significance of its deposition in the evolution of land plant cytokinesis and cell wall assembly in the context of plant’s migration to land.

**Figure 7:**
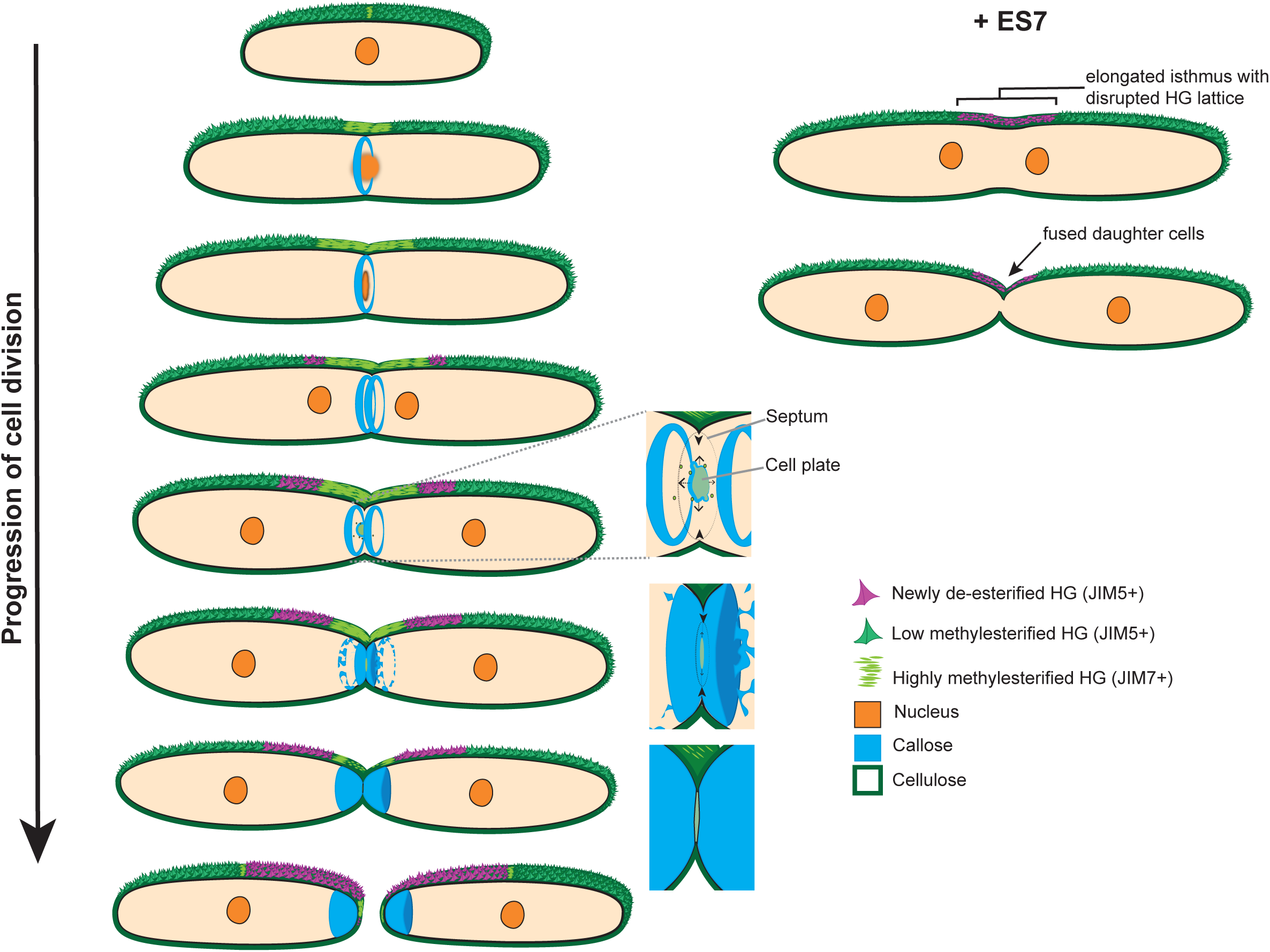
Model of cytokinesis in *P. margaritaceum*. Dividing *P. margaritaceum* cells first deposit callose in a single ring as early as metaphase. As cytokinesis progresses the single callose ring transforms into two rings prior to the appearance of a central callose punctum that we hypothesize marks the cell plate. Pectin deposition at the isthmus continues during this process, with lattice formation (JIM5+) following the secretion of methyl-esterified HG (JIM7+), de-esterification and calcium crosslinking. The callose rich (putative) cell plate then expands outward to meet the ingrowing septum. Once the cell plate meets the septum, cell wall formation continues with the deposition of cellulose and pectin to form two daughter cell poles. ES7 treatment disrupts the deposition of callose at the isthmus, which inhibits cytokinesis and causes elongation of the isthmus zone and fused cells. The lack of callose during cytokinesis subsequently disrupts the normal HG lattice formation.

## Materials and methods

### Growth and synchronization of cells

*Penium margaritaceum* (SKD-8 isolate, Skidmore College Algal Culture Collection) was maintained in liquid Woods Hole Medium (WHM) (Nichols 1973) supplemented with 1.5% (v/v) soil extract (Carolina Biological Supply, Burlington, NC, USA) as previously described by Domozych et al., 2007. Cells were maintained in a 30 mL cultures in vented falcon culture flasks (Fisher Scientific, Hampton, NH, USA). Cells were subcultured weekly by diluting 3 mL of 7 day-14 day log phase cultures to 27 ml of fresh WHM. *P. margaritaceum* cultures were grown in ambient temperature (∼20-22°C) in an LED-lighted cabinet with a photoperiod of 12 h. Cell synchronization was achieved within 4 days after subculture in new media.

*Spirogyra* sp. (Carolina Biological Supply) was maintained in 3N-BBM (Andersen, 2005; Bischoff and Bold, 1963; Starr and Zeikus, 1993) in ambient temperature and light conditions. New subcultures were made biweekly by adding several filaments to new media.

### ES7 treatment

Log phase cells were washed three times with fresh WHM and the pellet was resuspended in WHM supplemented with the indicated concentration of Endosidin 7 (ES7; ChemBridge Corporation, San Diego, CA, USA) of either 5µM, 7.5µM, 10µM or 50µM diluted in DMSO. DMSO control cells were cultured in WHM supplemented with an equal volume of DMSO alone. Cells were grown for 48 h in the respective media in the same light conditions as stated above prior to labeling and imaging.

For each treatment, three or more filaments of *Spirogyra* sp. were collected in a microcentrifuge tube with a sterile bacterial loop and washed three times with 3N-BBM by alternating light vortexing and pipetting. The filaments were added to fresh media supplemented with DMSO or 50 µM ES7 and allowed to grow for 48 h prior to labeling and imaging.

### Cell culture analysis

Cell culture absorbance at a wavelength of 600nm was measured every 24 hours with a Shimadzu spectrophotometer (UV-1700, Shimadzu, Kyoto, Japan), while WHM was used as control. ES7 phenotyping was carried out by observation of treated and non-treated cells under bright field (BF) microscopy. Isthmus zone length was measured using BF images of *P. margaritaceum* cells and the line tool in FIJI (Schindelin et al., 2012).

### Live cell labeling and analysis

ES7 and control treated *P. margaritaceum* cells were collected by centrifugation (2,000g, 1 min) and washed three times in WHM. Cells were resuspended in a 1:1000 dilution of SYTO9 (Invitrogen, Carlsbad, CA, USA) (in water) and vortexed for ∼30 s to label nuclei. Cells were then washed once with water and pellets were resuspended and incubated in 500 µL of a 1:40 (v/v) dilution of aniline blue fluorochrome (BioSupplies, Yagoona NSW 2199, Australia) in water. Cells were incubated in aniline blue fluorochrome for 10 min at room temperature in the dark. Cells were then mounted on slides for imaging. For labeling *Spirogyra* sp. filaments, the ES7 or DMSO media was removed by pipetting and the filaments were labeled with aniline blue fluorochrome as described above for *P. margaritaceum*.

### Live cell immunolabeling and analysis

Prior to immunolabeling, *P. margaritaceum* cells (treated and control) were washed three times in fresh WHM, then HG immunolabeling was carried out as previously described in Domozych et al. 2011 and Worden et al. 2015. Briefly, 3 mL of treated or control cells were collected by centrifugation, washed three times with fresh WHM and blocked with 1% (w/v) powdered milk (Signature select, Pleasanton, CA, USA) for 20 min. The cells were then washed and labeled with the respective primary antibody for 1.5 h and washed and blocked again prior to labeling with the secondary antibody at RT in the dark. During recovery experiments cells were first treated as described above, then washed six times with DMSO supplemented WHM, and grown for 2 days prior to relabeling with JIM5 antibodies. The cell wall directed antibodies were diluted in fresh WHM at the indicated concentrations: 1:10 dilution of rat-derived JIM5 directed against low methylesterified pectin (PlantProbes, University of Leeds, Leeds, England) and 1:8 dilution of JIM7 directed against highly methylesterified pectin (rat-derived, PlantProbes). Anti-rat Alexafluor488 (Invitrogen) diluted to 1:50 (in WHM) was used as the secondary antibody for both JIM5 and JIM7. After the 2 day recovery, anti-rat TRITC (Invitrogen) was used as the secondary antibody to distinguish new cell wall growth from previous growth.

### Fixed cell polysaccharide labeling

Cellulose was labeled with GFP-CBM3a (family 3 carbohydrate binding module; NZYtech, Lisboa, Portugal) following fixation with 1.5% PFA (PFA; Electron Microscopy Sciences, Hatfield, PA, USA) plus 0.5% glutaraldehyde (Ted Pella, Redding, CA, USA) at room temperature for 20 min. Cells were then incubated in pectate lyase for 20 min prior to washing and incubation with GFP-CBM3a for 90 min. Cells were then washed and counterstained with 1xDAPI prior to imaging. Callose labelling was performed by freeze shattering the cells and subsequent cell wall permeabilization as previously described in Wasteneys et al. 1997. Briefly, ES7-treated and control-treated cells were collected by centrifugation, washed three times with fresh WHM and fixed in 1.5% paraformaldehyde (Electron Microscopy Sciences) + 0.5% glutaraldehyde (Ted Pella) in 1x phosphate buffered saline buffer (PBS) for 30 min at room temperature. The cells where centrifuged at 2,000g and the resulting pellet was resuspended in ∼15µL, making a dense suspension that was sandwiched between two microscope slides and frozen by liquid nitrogen prior to shattering. Freeze shattered cells were washed into microcentrifuge tubes and subjected to a series of incubations and washes with permeabilizing buffers and then incubated with primary antibodies overnight at 4°C. Labeling was conducted as previously described in Ochs et al. 2014, with 1:1000 anti-callose primary antibody (mouse-derived, Biosupplies). Callose antibody-labeled cells were then incubated with 1:500 anti-mouse Alexa488 secondary antibody (Life Technologies) in the dark for 1.5 h prior to final washing. Cells were counterstained with 1x DAPI to visualize the nucleus and mounted on slides.

### Microscopy

All microscopy was conducted using a 710 Zeiss Axio inverted Observer.Z1 laser scanning confocal microscope (Carl Zeiss AG, Oberkochen, Germany). The following settings were used to image the corresponding dyes/antibodies: aniline blue fluorochrome (405nm excitation laser at 2% intensity, emission 410-502nm), SYTO9 (488nm excitation laser at 2% intensity, emission 500-567nm), GFP-CBM3a (488nm excitation laser at 10% intensity, emission 493-557nm), Alexa488 (488 excitation laser at 5% intensity, emission 493-630nm) used as secondary antibody for anti-callose, JIM5 and JIM7 experiments with the exception of the ES7 recovery experiments in which TRITC secondary antibodies were used (561nm excitation laser at 2% intensity, emission 566-601nm). All *P. margaritaceum* images were collected with Plan-Apo 40X (water, 1.1 NA) and 60X (oil, 1.4 NA) objectives. *Spirogyra* sp. images were collected with Plan-Apo 20X (air, NA) and 40X (water, 1.1 NA) objectives. Image analysis and figure preparation was conducted using Zen Black and Blue (Zeiss) imaging software, Adobe Photoshop and Illustrator. Surface rendering of fluorescent images was performed using Imaris (BitPlane, South Windsor, CT, USA).

### Phylogenetic analysis and tree construction

Previously characterized Arabidopsis thaliana GSL1-12 (Dong, 2004) sequences were used as queries for a BLAST search against representative genomes of different groups of Embryophyta, Charophyta, and Chlorophyta accessed at Phytozome 12 (phytozome.jgi.doe.gov) and the *Klebsormidium* genome project (<u>plantmorphogenesis.bio.titech.ac.jp</u>). The proteome database of *P. margaritaceum* was provided by Jiao et al. 2020, available at the Penium Genome Database (http://bioinfo.bti.cornell.edu/cgi-bin/Penium/home.cgi). GSL homologs showing over 50% amino acid similarity against the queries with e-values lower than 1^e-17^ were included in the analysis. Full-length GSL protein sequence alignment was performed using T-Coffee (Notredame et al., 2000), and used as the input for unrooted phylogenetic tree construction with Mega 10 (Kumar et al., 2018) using the maximum-likelihood method and 1000 bootstrap replication. Analysis was performed with default settings in MEGA 10 with the partial deletion site coverage set to 50%.

---

**Supplemental Figure 1: Cell wall growth and cell division recovers from ES7 in a majority of cells, revealing a callose-dependent cytokinesis checkpoint**.

ES7 [50] treated cells were first labeled with JIM5:FITC (green = initial cell wall) then allowed to recover prior to relabeling with JIM5:TRITC (magenta = new growth). Several JIM5 labeled HG patterns were apparent including those indicative of elongation and recently divided cells: (A) an elongating cell with new cell wall growth from the isthmus zone (magenta) prior to division; (B and C) exhibit cells that have recently divided with different levels of prior elongation; (D) recently divided cell (magenta tip) with isthmus zone elongation (central magenta signal). JIM5 relabeling revealed a population of cells (9.1%) that were unable to fully recover from ES7 treatment, with some cells exhibiting no new growth (E) and others capable of elongation prior to division and elongation at daughter isthmus zones (F). Scale bars = 20 µm.

**Supplemental Figure 2: Cellulose patterning during cytokinesis under callose inhibition**.

(A-C) In DMSO control cells cellulose labeling by GFP-CBM3a was observed in a single isthmus zone ring (A, arrowhead) and diffuse labeling over the entire cell that persisted throughout mitosis (B; n = 8). At the stage where daughter nuclei (blue) migrated to daughter isthmus zones, two cellulose rings were seen at the parental isthmus zone (C, arrowheads; n = 6). (D-H) In ES7-treated cells, normal cellulose ring patterning was observed in interphase (D, arrowhead; n = 2) and immediately following daughter nuclei migration (E, arrowhead; n = 6). In addition, ES7-treated cells exhibited ectopic cellulose rings (n = 3) that varied from multiple (G and H, arrowheads) to improperly localized rings (F, arrowheads). Scale bars = 20 µm. Images are max intensity projections.

**Supplemental Figure 3. Domain analysis of glucan synthase-like enzymes**.

Domains recognized by NCBI Conserved Domain Analysis (Shennan Lu et al, 2020) is indicated by colored boxes and unknown regions are indicated by white boxes. The length of the boxes is proportional to number of amino acid residues. 15 putative full-length *P. margaritaceum* protein sequences aligned to AtGSL2 is indicated below AtGSL2, with sequences over 50% amino acid similarity to AtGSL2 drawn with boxes. The black bars indicate the length of amino acid similarity coverage and grey lines indicate no similarity of the corresponding region.

## Acknowledgements

We would like to thank Drs Bo Liu and Judy Callis for their useful suggestions. We thank Rosalie Sinclair and Oliver Betz for setting up the growth incubators and for their useful comments.

## List of symbols and abbreviations

ES7: Endosidin7
HG: homogalacturonan
WHM: Wood’s Hole Medium
3N-BBM: triple nitrogen Bold’s Basal Medium

